# Membrane interactions of mitochondrial lipid transfer proteins

**DOI:** 10.1101/2022.04.05.487160

**Authors:** Fereshteh Sadeqi, Kai Stroh, Marian Vache, Dietmar Riedel, Andreas Janshoff, Herre Jelger Risselada, Michael Meinecke

## Abstract

The mitochondrial inner membrane is an integral part of the cellular lipid biosynthesis network. Intramitochondrial lipid transfer shuttles specific lipid species between the two mitochondrial membranes. This pathway is facilitated by designated protein complexes in the intermembrane space. A hetero-dimeric complex of Ups1 and Mdm35 has been identified in yeast to transfer phosphatidic acid from the outer to the inner membrane. Here, we define pH and lipid charge dependency of Ups1 membrane interaction. We show that Ups1 interacts with membranes in a membrane curvature dependent manner. Ups1 predominantly binds to membrane domains of positive curvature. Desorption of phosphatidic acid from positively curved membranes is energetically favorable over flat or negatively curved membranes. Hence, our results demonstrate that Ups1 specifically binds to membrane regions where extraction and insertion of lipids is enhanced.

## Introduction

Eucaryotic cells display a high degree of compartmentalization characterized by the appearance of various membrane enclosed organelles. Organellar membranes function on the one hand as barriers to create specialized reaction rooms. On the other hand, vital biochemical pathways, lipid biosynthesis or ATP production to name just two, take place at or within membranes themselves. Consequently, the composition of organellar membranes is highly diverse with respect to their protein and lipid content and is adjusted to its physiological function (Meer et al., 2008). Besides membranes of the endoplasmic reticulum (ER) the mitochondrial inner membrane (IM) is an integral part of the cellular lipid biosynthesis network. Here, among others, the mitochondrial signature lipid cardiolipin (CL) is produced in a multi-step reaction (Osman et al., 2011). To this end phosphatidic acid (PA) has to be imported from the ER and transferred from the mitochondrial outer membrane (OM) to the IM. For this non-vesicular transport of PA, a specific heterodimeric protein complex comprised of yeast Ups1 (PRELID1 in mammals) and Mdm35 (TRIAP1) has been identified (Osman et al., 2009; Tamura et al., 2009). Ups1/Mdm35 is located in the intermembrane space (IMS) of mitochondria. *In vitro* studies showed that Ups1 binds to membranes and can directly transfer PA between membranes upon dynamic assembly with Mdm35 (Connerth et al., 2012). Crystal structures of Ups1/Mdm35 in the absence and presence of PA were obtained showing that one lipid molecule per heterodimeric complex is bound (Watanabe et al., 2015; Yu et al., 2015). These results give rise to the following model. Ups1 binds to the IMS surface of the OM most likely under dissociation of Mdm35. A PA molecule is extracted from the OM and Ups1 dissociates under reassembly with Mdm35. PA is shuttled through the aqueous IMS. Ups1 binds to the IMS surface of the IM under dissociation of Mdm35 and releases the lipid into the IM (Tamura et al., 2020).

One of the major questions that arise from this model is how the energy barrier is overcome to break lipid-lipid interactions extract a lipid from one membrane and insert it into a different one. To tackle this question, we used a combination of *in vitro* and *in silico* experiments to quantitatively analyze Ups1/Mdm35 membrane interaction.

## Results & Discussion

### Ups1 binds to membranes in a charge and pH dependent manner

To study protein-membrane interaction we first recombinantly expressed and purified yeast Ups1/Mdm35 as a heterodimeric complex from *E. coli* cells (Figure 1 A). We next used liposomes flotations assays to follow membrane binding of Ups1/Mdm35. Large unilamellar vesicles (LUVs) with a diameter of 200 nm, roughly resembling the IM composition (55 % L-α-phosphatidylcholine (PC), 20 % L-α-phosphatidylethanolamine (PE), 10 % CL, 15 % PA (L-α-phosphatidic acid)), were incubated with Ups1/Mdm35. Liposomes were separated from unbound protein by Histodenz gradient centrifugation, which was subsequently fractionated and analysed by SDS PAGE (Figure 1 B). Liposomes clearly floatated to upper fractions of the gradient (Figure 1B). As shown before (Connerth *et al*., 2012; Lu et al., 2020) Ups1 bound to and floated with liposomes in a pH dependent manner while Mdm35 predominantly stayed in the unbound bottom part of the gradient (Figure 1B). At pH 5.5 about 95 % of Ups1 bound to membranes while that value dropped to less than 40 % at pH 7.4. To determine binding constants of Ups1 to membranes we performed reflectrometric interference spectroscopy (RIfS) measurements using the same lipid composition (Figure C). At a pH of 5.5 we measured a dissociation constant of K_D_ = 0.38 ± 0.5 μM, whereas at pH 7.4 binding was not resolvable by RIfS. When using membranes only consisting of PC and PE no binding was observed regardless of the pH (data not shown).

**Figure 1:**
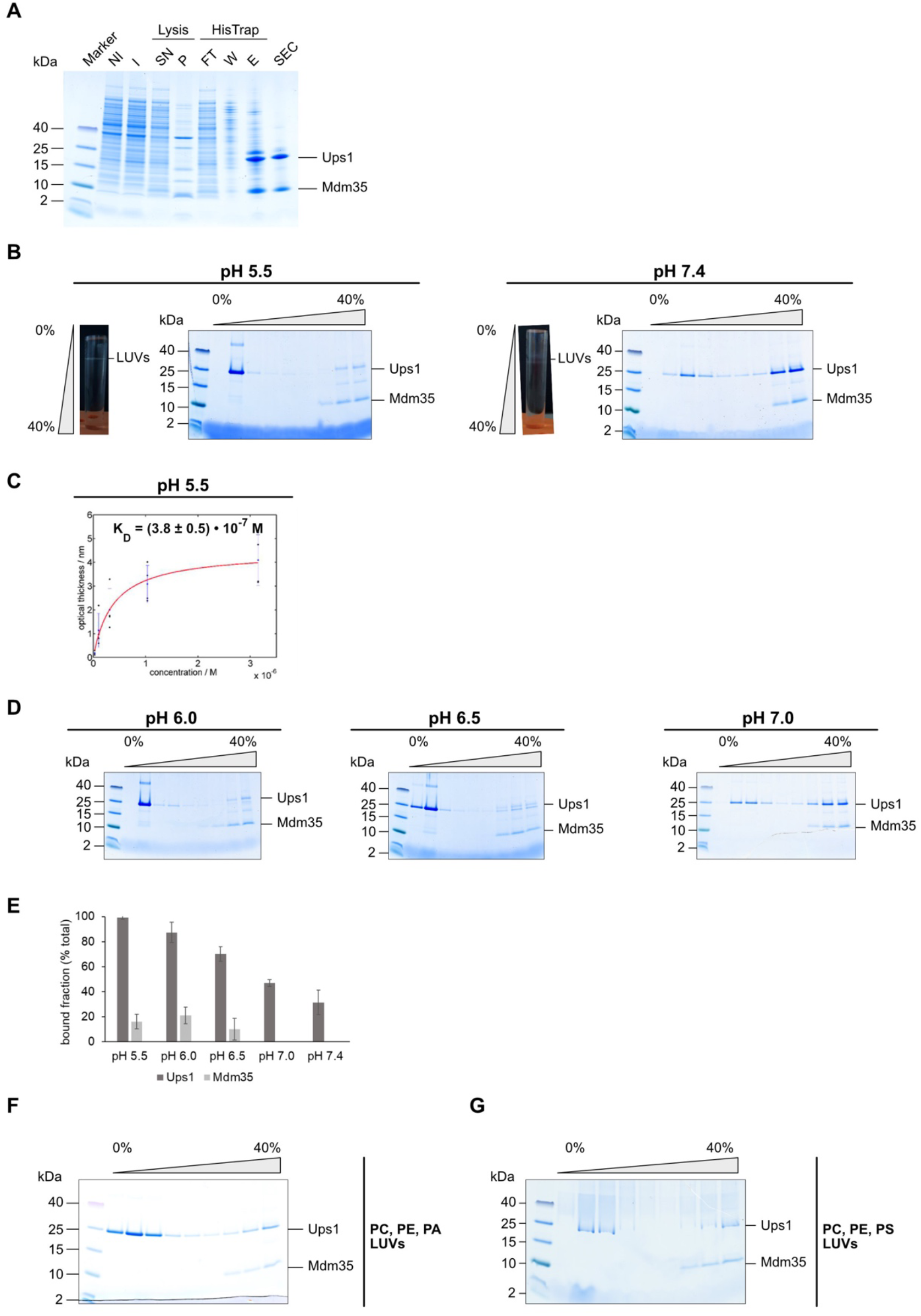
Ups1 binds to membranes in a charge and pH dependent manner. (**A**) Expression and purification of recombinant Ups1/Mdm35 complex analyzed by SDS-PAGE and colloidal Coomassie staining. NI = non induced, I = Induced, SN = supernatant, P= pellet, FT = flowthrough, W = wash, E = elution, SEC = size-exclusion chromatography (**B**) Flotation of LUVs (55%PC/20%PE/10%CL/15%PA) with Ups1/Mdm35 in a non-continuous Histodenz density gradient at pH 5.5 and pH 7.4. Samples were fractionated after flotation and analyzed by SDS-PAGE and colloidal Coomassie staining. (**C**) Dissociation constant (*K*_D_) for Ups1/Mdm35 to membranes containing 55%PC/20%PE/10%CL/15%PA. The *K*_D_ was obtained by a Langmuir fit of the optical thickness. (**D**) Fractionation of flotated Ups1/Mdm35 with LUVs (55%PC/20%PE/10%CL/15%PA) at pH 6.0, pH 6.5 and pH 7.0 analyzed by SDS-PAGE and colloidal Coomassie staining. (**E**) Quantification of the bound fraction (% total) seen on the SDS gels of pH-dependent flotation assays. For each of the proteins Ups1 and Mdm35, the sum of band intensities in all nine fractions was set to 100 %. Quantification was done for three independent experiments and values for bound fraction (% total) are shown as the mean. Error bars represents the SD. (**F**) Fractionation of flotated Ups1/Mdm35 with LUVs (50%PC/20%PE/30%PA) and (**G**) LUVs (50%/20%/30%PS) analyzed by SDS-PAGE and colloidal Coomassie staining.

Due to the proton pumping activity of OXPHOS complexes within the IM, the pH of the IMS is lower than in the cytosol and in the matrix and was measured to be around 6.8 (Porcelli et al., 2005; Santo-Domingo and Demaurex, 2012). If there is pH gradient within the IMS is a matter of debate. Nonetheless, it is tempting to speculate that at the IMS surface of the IM, where protons appear after pumping, the pH is lower than at the IMS surface of the OM, which contains several water-filled β-barrel porins. We therefore asked whether we see differences in Ups1 membrane binding in this physiological range. Indeed, flotations at various pH values showed a turning point of predominant bound to unbound protein fraction between pH 6.5 and pH 7.0 (Figure 1D). If this switch plays a role in regulating the directionality of lipid transfer will be an exciting question for future studies.

Ups1 membrane binding was shown in the past to depend on negatively charged lipids. To investigate whether this is a pure electrostatic effect or if it includes some degree of lipid specificity, we performed flotation experiments with LUVs comprised of PC and PE spiked with either 15 % CL, 30 % PA or 30 % PS. Binding experiments showed that Ups1 efficiently binds to all LUVs though CL containing vesicles showed the strongest binding (Figure 1F & G).

### Ups1 has a membrane curvature sensing activity

Membrane binding of Ups1 depends, at least partially, on the hydrophobic α2-loop (Lu *et al*., 2020; Miliara et al., 2019). The insertion of hydrophobic protein domains into membranes was shown many times to lead to membrane remodeling (Steinem and Meinecke, 2020; Tarasenko and Meinecke, 2021). Hence, we asked whether binding of Ups1 to membranes leads to the induction of membrane curvature. LUVs in the absence and presence were visualized via electron microscopy. Whereas in the absence of protein almost all visualized vesicles appeared spherical, after protein addition about 35 % of the vesicles showed deformed morphology (Figure 2A & B). It should be noted that the membrane remodeling effects of Ups1 appeared less pronounced than for proteins involved in membrane trafficking or, for example, mitochondrial proteins that deform the inner membrane at crista junctions (Farsad et al., 2001; Henne et al., 2010; Hessenberger et al., 2017; Tarasenko et al., 2017).

**Figure 2:**
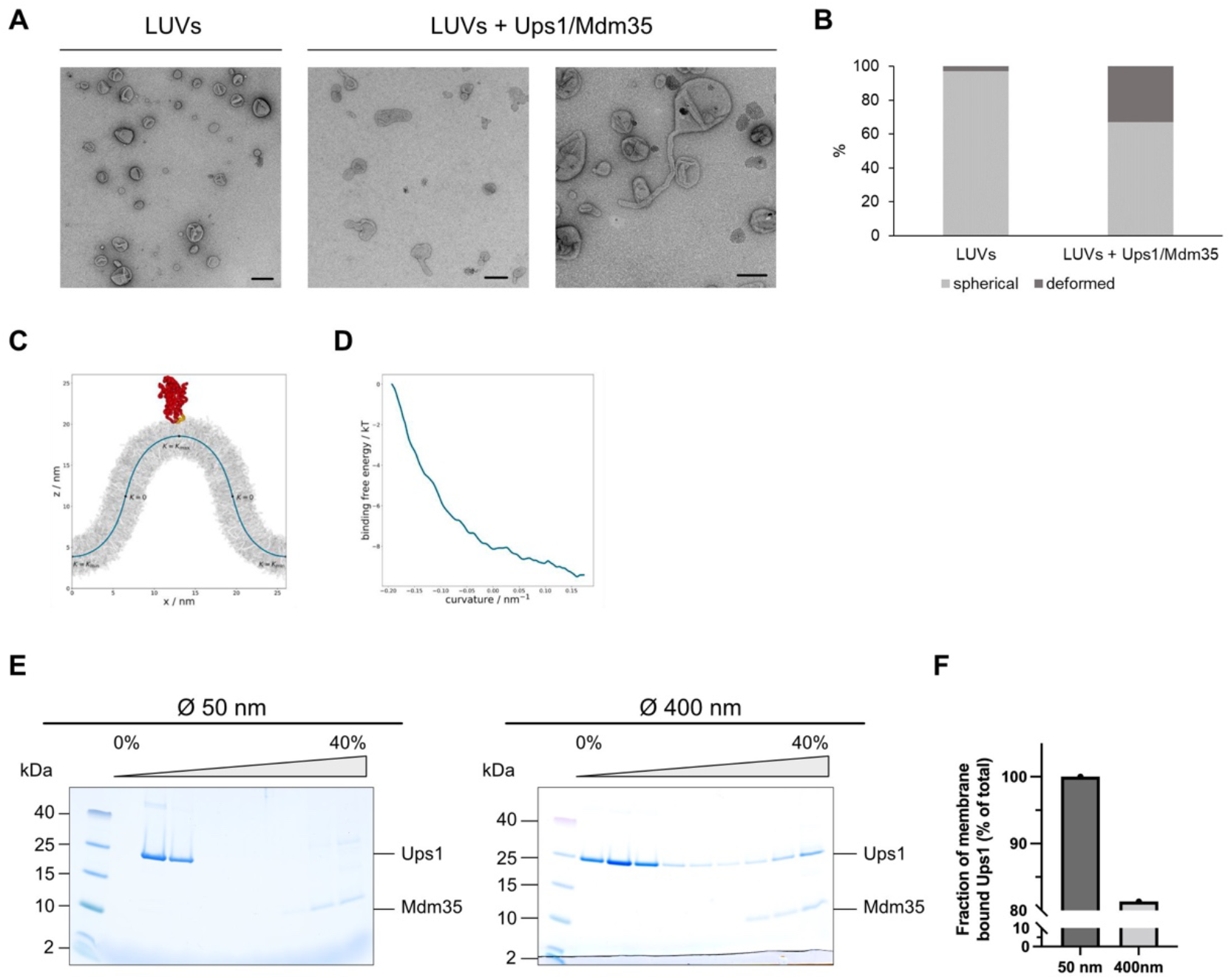
Ups1 has a membrane curvature sensing activity. (**A**) Electron micrographs of liposomes (LUVs) in the absence and presence of the protein complex Ups1/Mdm35. (**B**) Quantification of spherical and deformed shapes in control LUVs and in LUVs after incubation with Ups1/Mdm35. A minimum of 200 membranous structures were counted per condition. Scale bar corresponds to 100 nm. (**C**) The Ups1/Mdm35 complex (in red, hydrophobic loop in yellow) on a buckled membrane and analytical membrane shape description. (**D**) Relative binding free energy as a function of membrane curvature. (**E**) Fractionation of flotated Ups1/Mdm35 with LUVs (50%PC/20%PE/30%PA) of 50 nm and 400 nm size. (**F**) Quantification of co-floated Ups1 after incubation with 50 nm or 400 nm sized liposomes, respectively.

Another effect of hydrophobic or amphipathic protein domains is that they act as membrane curvature sensors. Such domains sense lipid packing defects accompanying deformed membranes. With this activity proteins can be guided to predominantly bind to membrane regions of a certain curvature (Antonny, 2011). To analyze whether Ups1 displays a membrane curvature sensing activity we first used an *in silico* approach to directly resolve the relative partitioning free energy of proteins as a direct function of membrane curvature, i.e., a curvature sensing profile, within (coarse-grained) molecular dynamics simulations (Stroh and Risselada, 2021). For a membrane curvature ranging from −0.188 nm^−1^ (K_min_) to +0.188 nm^−1^ (K_max_) we obtained a change in the binding (partitioning) free energy of ΔF = 9.2 kbT ± 1.5 kbT (95 % confidence interval), with Ups1 preferring partitioning to regions of positive membrane curvature (Figure 2C & D). We next aimed to show this membrane curvature sensing activity by biochemical means using the liposome co-flotation assay. Vesicles with diameters of 50 nm (high positive curvature) and 400 nm (low positive curvature) were constructed. As we could show before that CL containing LUVs showed extremely strong binding of Ups1, which might mask a curvature sensing activity, we used vesicles comprised of PC, PE and PA at pH 5.5. SDS PAGE analysis of the different gradient fractions clearly showed curvature dependent differences with higher degrees of Ups1 binding to smaller liposomes, i.e. to membranes with high degrees of positive curvature (Figure 2E). Both *in silico* and *in vitro* experiments thus show preferential binding of Ups1 to membrane regions of positive curvature.

What would the physiological advantage of protein-membrane interaction at regions of positive curvature be? To answer that question, we hypothesized that extraction and insertion of a PA molecule should be energetically more favorable at regions displaying lipid packing defects, i.e. positive membrane curvature. We used the same buckled membrane as before but now in the absence of protein to determine free energy landscapes for three major lipid species in this membrane (Figure 3). In this case no accelerated sampling is necessary and the potential of mean force (PMF) can be constructed directly from the lipid distributions along the reaction coordinate.

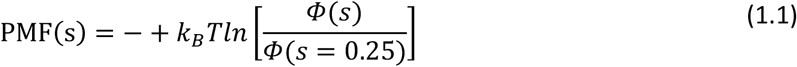

with the Boltzmann constant *k_B_*, the temperature *T*, the arc length parameter *s*, and the lipid distribution ϕ(*s*). Since the end state of an extraction process of a lipid (into the binding pocket of Ups1 or water) would be the same, regardless of the membrane region the lipid is extracted from. Relative free energy differences of extraction can be read off directly from these free energy landscapes. An increase of free energy of lipids inside the membrane results in a lowered amount of work for extraction. We found that for PA and PE the energy cost of desorption is decreased for positive curvature, and increased in negatively curved regions. This is in line with earlier results showing an enhanced rate of lipid extraction at regions of high positive membrane curvature (Collado et al., 2019). The curvature effect is reversed for PC. Compared to PC, PE and PA both have smaller headgroups and the positively curved region presents a less optimal environment for them. For PA charge repulsion most likely plays a role, as the curvature effect is smaller compared to PE, while it has an even smaller headgroup than PE.

**Figure 3:**
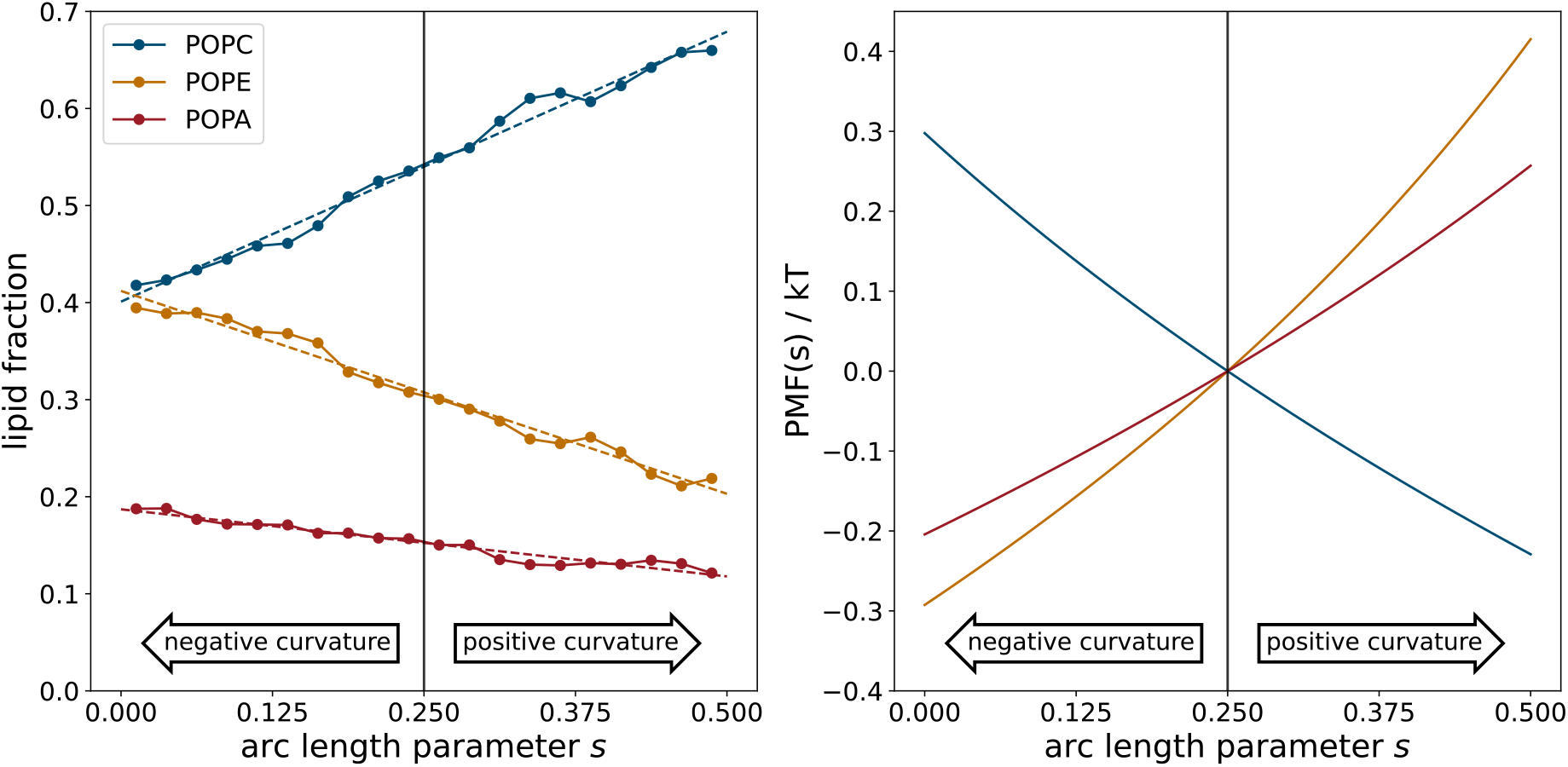
Membrane curvature dependent lipid extraction. (Left) Lipid distributions along the arc length parameter s. The membrane is at s = 0:25. Errors from bootstrapping are smaller than the symbols. Dashed lines represent the mean of linear fits to every resampled lipid distribution. (Right) Free energy landscapes of each lipid species along the membrane arc constructed from the mean of linear fits to the lipid distributions.

One should note that the effects described above are composition dependent, as changes to composition also change the partitioning behavior (Boyd et al., 2017; Elias-Wolff et al., 2019). Together our results demonstrate that Ups1 shows preferential binding to membrane domains of positive curvature where, in turn, the energetic cost to extract a lipid is lowered. As we also found a mild membrane curvature inducing activity, lipid transfer proteins might very well be able to induce lipid packing defects and thus not only bind to but even create a microdomain that facilitates lipid transfer. These results offer a molecular explanation of how the energy barrier to extract and insert a lipid from and into a membrane is affected by lipid transfer molecules themselves. Membrane curvature at the level of the IM is well described (Barbot and Meinecke, 2016; Mukherjee et al., 2021). High degrees of positive curvature are found at crista junctions where the MICOS complex subunits Mic10 and Mic60 induce curvature (Barbot et al., 2015; Hessenberger *et al*., 2017; Tarasenko *et al*., 2017). Membrane remodeling in the OM is less well understood than in the IM. Nonetheless, also OM proteins with the ability to deform membranes have been found (Lackner et al., 2009) and structural investigations of OM residing proteins imply that, at least, micro-domains exhibiting membrane curvature might be a more wide-spread phenomenon (Bausewein et al., 2017; Takeda et al., 2021). It is hence likely that in both mitochondrial membranes domains with lipid packing defects exist. Interestingly, the curvature inducing Mic60 is also part of a membrane bridging complex that, together with Sam50 in the OM, physically links IM and OM (Sastri et al., 2017; Tang et al., 2019). Membrane-membrane contact sites have been found before to facilitate lipid transfer pathways (Egea, 2021). Consequently, we hypothesize that proteins like Mic60 not only provide the curvature to recruit lipid transfer proteins but at the same time bring IM and OM in close contact, which additionally could enhance lipid transfer rates. If and how these proteins work in cooperativity will be formidable questions for future studies.

## Methods

### Recombinant protein expression and purification

Expression and purification of Ups1/Mdm35 was performed according to previously published protocols (Connerth *et al*., 2012). The coding sequence for *UPS1* carrying an N-terminal His-tag, and *MDM35* from *Saccharomyces Cerevisiae* were cloned into the pET-Duet-1 (Merck) vector and co-expressed in Origami B(DE3)pLysS *E.coli* cells (Merck) in the presence of 1 mM IPTG at 30°C for 3h. The cells were harvested and resuspended in lysis buffer [250 mM NaCL, complete protease inhibitor cocktail without EDTA (Roche), 1 mM PMSF, 50 mM Tris/HCl pH 8.0]. Cell lysis was performed using an EmulsiFlex-C5 (Avestin). The lysate was centrifuged and the supernatant was filtered through a 0.45 μm filter and subjected to a His-tag affinity chromatography using a 5 mL HisTrapFF (GE Healthcare) and the ÄKTAprime plus system (GE Healthcare). The bound proteins were eluted with 250 mM NaCL, 500 mM Imidazole, 50 mM Tris/HCl, pH 8.0 and the peak fractions were collected and concentrated using an Amicon filter (Merck Milipore). The sample was further purified via size-exclusion chromatography (SEC) with a HiLoad 16/600 75 pg (GE Healthcare) column in buffer containing 150 mM NaCL, 50 mM Tris/HCl, pH 7.4. After collecting the peak fractions, the sample was stored at −80°C.

### Liposome preparation

Liposome preparation was done as described in detail before (Denkert et al., 2017). The lipids L-α-Phosphatidylcholine (PC), L-α-Phosphatidylethanolamine (PE), L-α-Phosphatidic acid (PA), L-α-Phosphatidylserine (PS) and cardiolipin (CL) were purchased from Avanti Polar Lipids. Lipids were mixed in the desired molar ratios: PC/PE/PA (50:20:30), PC/PE/PA (65:20:15), PC/PE/CL/PA (55:20:10:15), PC/PE/CL (65:20:15); PC/PE/PS (65:20:15); PC/PE/PS (50:20:30). The lipid mixtures were dried shortly under a stream of nitrogen gas, followed by an incubation in a desiccator for at least 2 h. The dried lipids were hydrated in the corresponding liposome buffer at desired pH (150 mM NaCl, 10 mM MES/NaOH, pH 5.5); (150 mM NaCl, 10 mM MES/NaOH, pH 6.0); (150 mM NaCl, 10 mM MES/NaOH, pH 6.5); (150 mM NaCl, 50 mM Tris/HCl, pH 7.0); (150 mM NaCl, 50 mM Tris/HCl, pH 7.4). LUVs were prepared by performing freeze and thaw cycles while SUVs were generated by an additional step of sonification after the freeze and thaw cycles. The liposomes were subjected to extrusion through polycarbonate membranes (Whatman) with a desired pore size (50 nm, 200 nm or 400 nm). For RIfS measurements, the lipids PC/PE/CL/PA were mixed in the following molar ratio 55:20:10:15. After drying, the generation of SUVs was performed by swelling the lipid films in buffer (150 mM NaCl, 10 mM MES/NaOH, pH 5.5) followed by sonication (Sonoplus HD 2070, Bandelin).

### Liposome flotation

For the flotation assay 10 μM of the Ups1/Mdm35 complex was incubated (for 30 min at room temperature) with LUVs or SUVs (5 mM of total lipids) in a total reaction volume of 100 μL in the corresponding liposome buffer. The sample was mixed with 700 μL of 40 % Histodenz and transferred into the bottom of an ultracentrifuge tube. The mixture was overlaid with 20 %, 10 % and 5 % Histodenz (à 900 μL) as well as the corresponding liposome buffer on top, creating a non-continuous Histodenz gradient from high density to low density (40%/20%/10%5%/0%). The sample was subjected to ultracentrifugation at 55000 rpm at 4 °C for 1 h. Nine fractions à 500 μL were taken for precipitation with 10 % TCA and subsequently prepared for SDS-PAGE analysis.

### Reflectometric interference spectroscopy (RIfS)

Wafers: 5000 nm Silicon oxide silicon wafers, Active Business Company, Brunnthal, Germany Other materials used can be found in Witt *et al*. (Witt *et al*. 2020).

#### Reflectometric interference spectroscopy (RIfS) measurements

Reflectometric interference spectroscopy experiments were performed with a home-built setup, consisting of a tungsten halogen light source (LS-1, Ocean Optics, Dunedin, Florida, USA), a flow chamber transparent for visible light which firmly covers a silicon wafer with 5 μm silicon oxide layer (Active Business Company, Brunnthal, Germany) and a spectrometer (Nanocalc-2000-UV/VIS, Ocean Optics). The detailed setup and data analysis to obtain values of the optical thickness is described elsewhere (Stephan 2014).

The flow chamber was first purged with buffer containing 10mM MES, 150 mM NaCL, at pH 5.5. To create a supported membrane, the PC/PE/CL/PA (55:20:10:15) containing SUV dispersion was flushed through the system until reaching a stable plateau of the optical thickness. After rinsing of the supported membrane with buffer, the system was put into a solution of Ups1/Mdm35. In order to obtain a Langmuir isotherm, the experiment was conducted with increasing concentration of Ups1/Mdm35. Lastly, the membrane was rinsed with buffer. To analyse the RIfS data, the obtained optical thicknesses were plotted against the concentration and a Langmuir isotherm was fitted to the data according to eq. 1.

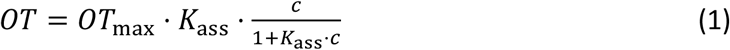

With *OT* being the optical thickness, *OT*_max_ the equilibrium optical thickness at the maximum concentration of protein, *c* the protein concentration and *K*_ass_ the thermodynamic association constant. The thermodynamic dissociation constant (*K*_d_) is the inverse of the association constant. Calculation of the errors of the fit of the means were obtained using the MATLAB (MATLAB R2014a (MathWorks, Natick, MA) function confint.

### Electron Microscopy of LUVs

LUVs in the presence or absence of protein were bound to a glow discharged carbon foil covered 400 mesh grid and stained with 1% uranyl acetate solution as described before (Vasic et al., 2020). Electron microscopic imaging and visualization of the sample was performed at room temperature with a Talos L120C transmission electron microscope (ThermoFischer Scientific, Eindhoven, and The Netherlands).

### In silico curvature sensing

Coarse-grained molecular dynamics simulations were performed with GROMACS 2020.3 (Abraham et al., 2015) using the Martini 2.2 force field (de Jong et al., 2013; Monticelli et al., 2008). The atomistic protein structure (PDB 4XHR, (Yu *et al*., 2015)) was translated to a coarsegrained description using the martinize Python script (de Jong et al., 2013). Protein secondary structure was determined employing DSSP (Kabsch and Sander, 1983; Touw et al., 2015) and overall protein structure was stabilized via elastic band with a force constant of 500 kJ/(mol*nm^2^).

Initial configurations of the protein membrane system were generated with the python script insane (Wassenaar et al., 2015). 674 lipids per leaflet (371 POPC, 202 POPE, 101 POPA) were put in the x-y plane of a 40 nm * 10 nm * 20 nm simulation box. The protien complex was placed approcimately 1 nm above the upper leaflet. The system was solvated with standard Martini water and a 0.15 M NaCl concentration. Steepest-descent energy minimization and initial equilibration (10 ns NVT and 50 ns NPT) were followed by compressing the system in x-direction by applying a pressure of 3 bar in x-direction while allowing the system to expand in z-direction only (Berendsen barostat (τ_p_=12.0 ps, compressibility of 3 * 10^−4^ bar^−1^)(Berendsen et al., 1984)). From the compression trajectory a frame close to the chosen compression level was selected. The selected frame was then used as a starting configuration of a long equilibration phase that was 5 microseconds long (Parrinello-Rahman barostat (τ_p_=12.0 ps, compressibility of 3 * 10^−4^ bar^−1^)(Parrinello and Rahman, 1981)). This long run is necessary to allow curvature-induced lipid partitioning to finalize. For the umbralla sampling the buckled membrane was fixed to its analytical shape by position restraints on the headgroups of lower leaflet lipids with a small force constant (10 kJ mol^−1^ nm^−2^). Influence on upper leaflet dynamics is minimal with these restraints. To generate starting configurations for the umbrella sampling, the protein complex was pulled along the membrane in x-direction. 60 windows equidistant in arc length parameter s, between s = 0 and s = 0.5 were used. Umbrella sampling runs were 1.05 μs long, with the first 0.05 μs for equilibration. Harmonic potentials with a force constant k = 100 kJ mol^−1^ nm^−2^ were used to restraint the hydrophobic loop to a defined position along the reaction coordinate. Umbrella sampling simulations were performed in the NVT ensemble. All simulations were coupled to a constant heat bath using the velocity rescale algorithm (Bussi et al., 2007).

Lipid concentrations along the arc length parameter s were extracted from a 2 μs simulation in the NVT ensemble. System setup was as described above, except the protein complex was not present in this simulation. Since the membrane shape is mirror symmetric around s = 0.5, lipid distributions are avarged over both sides and presented from s = 0 to s = 0.5. Potentials of mean force were constructed from linear fits to the lipid distributions. Errors were estimated via bootstrapping (Newman, 1999). Each lipid distribution was resampled 1000 times.

## Acknowledgments

This work was funded by the DFG FOR2848 (P05 to FS and MM, Z01 to DR). The authors are grateful for the computational resources provided by the Jülich Supercomputing Center and the HLRN Berlin / Göttingen.

## Notes

### Competing Interest Statement

The authors have declared no competing interest.

